# Biofilm matrix proteome of clinical strain of *P. aeruginosa*

**DOI:** 10.1101/2021.12.21.473640

**Authors:** Daria A. Egorova, Andrey I. Solovyev, Nikita B. Polyakov, Ksenya V. Danilova, Anastasya A. Scherbakova, Ivan N. Kravtsov, Maria A. Dmitrieva, Valentina S. Rykova, Irina L. Tutykhina, Yulia M. Romanova, Alexander L. Gintsburg

## Abstract

Extracellular matrix plays a pivotal role in biofilm biology and proposed as a potential target for therapeutics development. As matrix is responsible for some extracellular functions and influence bacterial cytotoxicity against eukaryotic cells, it must have unique protein composition. *P. aeruginosa* is one of the most important pathogens with emerging antibiotic resistance, but only a few studies were devoted to matrix proteomes and there are no studies describing matrix proteome for any clinical isolates. Here we report the first biofilm matrix proteome of *P. aeruginosa* isolated from bronchoalveolar lavage of patient in intensive care unit. We have identified the largest number of proteins in the matrix among all published studies devoted to *P. aeruginosa* biofilms. Comparison of matrix proteome with proteome from embedded cells let us to identify several enriched bioprocess groups. Bioprocess groups with the largest number of overrepresented in matrix proteins were oxidation-reduction processes, proteolysis, and transmembrane transport. The top three represented in matrix bioprocesses concerning the size of the GO annotated database were cell redox homeostasis, nucleoside metabolism, and fatty acid synthesis. Finally, we discuss the obtained data in a prism of antibiofilm therapeutics development.

## 1. Introduction

Biofilms are the most common lifestyle of microorganisms, including both pathogenic and environmental bacterial species. From a clinical perspective, biofilm cause difficult-to-treat recurrent diseases. Microbial aggregates with tolerance to host defense mechanisms and antimicrobials are found at the site of infection. Currently, there is an urgent need to discover new targets and strategies to overcome the tolerance for the effective treatment of biofilm-associated infections.

The key feature of biofilms is an extracellular matrix that covers all members of biofilm and creates a microenvironment for communication, protects against different threats, provides an opportunity for spatial organization and functional diversification within the community. The matrix comprises a broad range of biopolymers, metabolites, and signal molecules. Also, it may include organized compartments like outer membrane vehicles (OMVs). To stress the idea of a rich and complex matrix organization, Karygianni at al. have proposed the term «matrixome» [1]. The sources of proteins in the biofilm matrix might be active secretion, passive leakage from cells, and entrapped into the matrix from the environment molecules [2,3]. Proteins in the matrix have a structural role in maintaining biofilm organization, play a role as a protective barrier, creates microenvironment with limited diffusion. The barrier role is somewhere similar to structural function but also includes hydrophobic features, charge, and so on, rather than just being mechanically stable structure. Moreover, matrix proteins may bind antimicrobial molecules and reduce their diffusion and effect on bacterial cells, i.e. extracellular ribosomal proteins bind antibiotics and prevent their penetration in bacterial cells. Other functions of matrix proteins rely on their enzymatic activity and include degradation of biopolymers, participation in biochemical processes, and signaling function.

Biofilms are dynamic communities. Lifecycle can be divided into several major stages: attachment, maturation, and dispersion. During the maturation stage, the matrix may accumulate virulence factors which then come out together with dispersed bacteria. During chronic infections, the dispersion stage is associated with the recurrence of symptomatic infections and colonization of new sites in the body.

Importantly, bacteria inside biofilm have diverse phenotypes, so one produces some molecules in extracellular space while non-producers consume these public goods. Matrix as a compartment is also an example of public goods. A part of bacterial population, especially inside multispecies biofilms, may not have tolerance or specific resistance mechanisms, but due to matrix still be irresponsible for the treatment.

Targeting of matrix proteins as a strategy to combat infections may act in the following ways: (1) destabilization/disruption of biofilm matrix to improve the action of host defense and/or antibiotic therapy; (2) targeting of extracellular biochemical processes to reduce overall biofilm success in survival and virulence; (3) targeting of extracellular virulence factors associated with biofilm matrix to decrease virulence in a case of mass dispersion.

Despite the accepted idea of a pivotal role of matrix in bacterial biofilms, protein composition of the matrix remains poorly discovered. Moreover, for the well-studied and clinically important biofilm-forming bacteria *P. aeruginosa* there are only a few studies devoted to matrix proteome of reference strains (mainly PAO1, Table 1), while proteomics of the whole biofilm is better described [4,5].

**Table 1.**
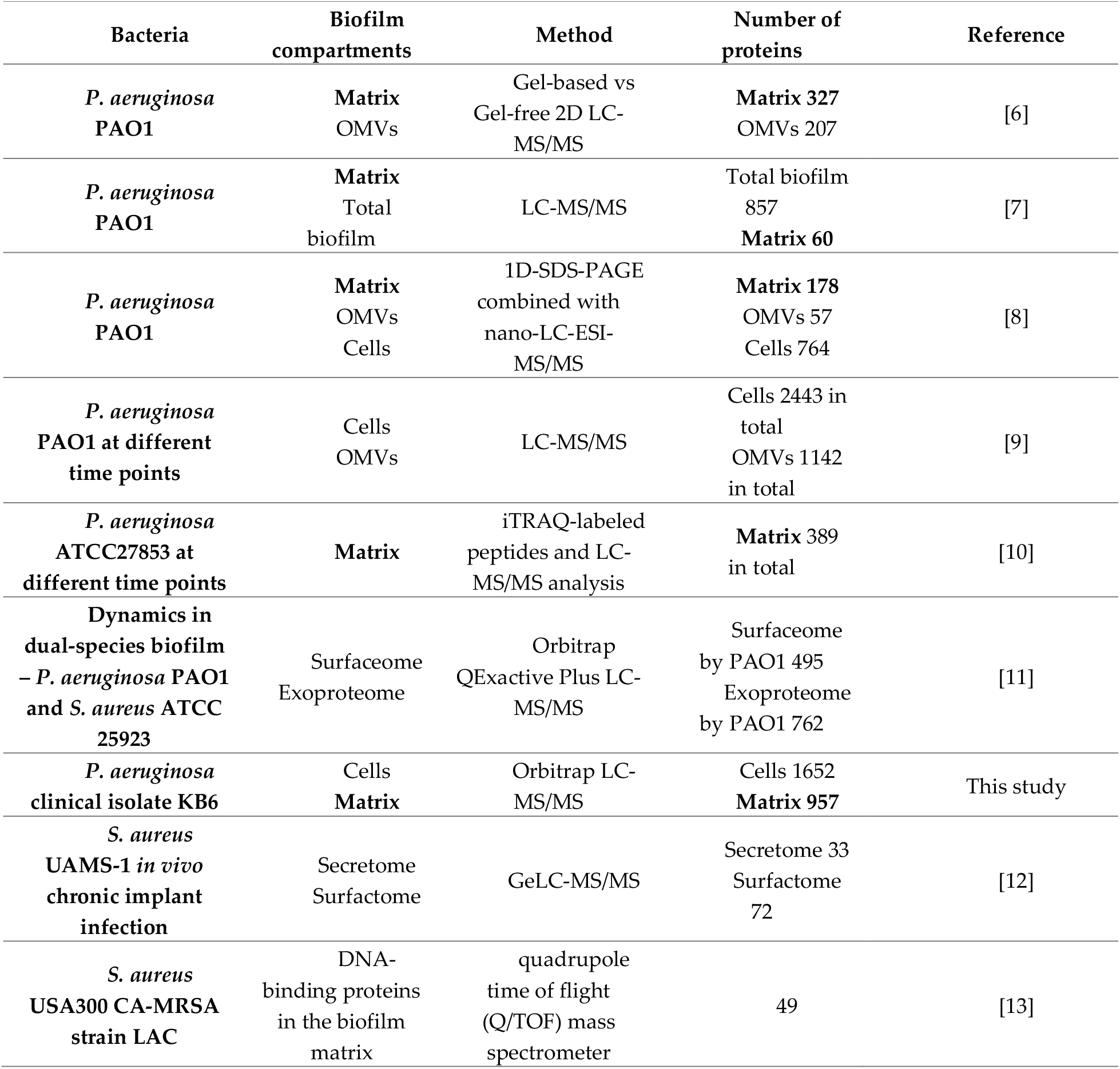
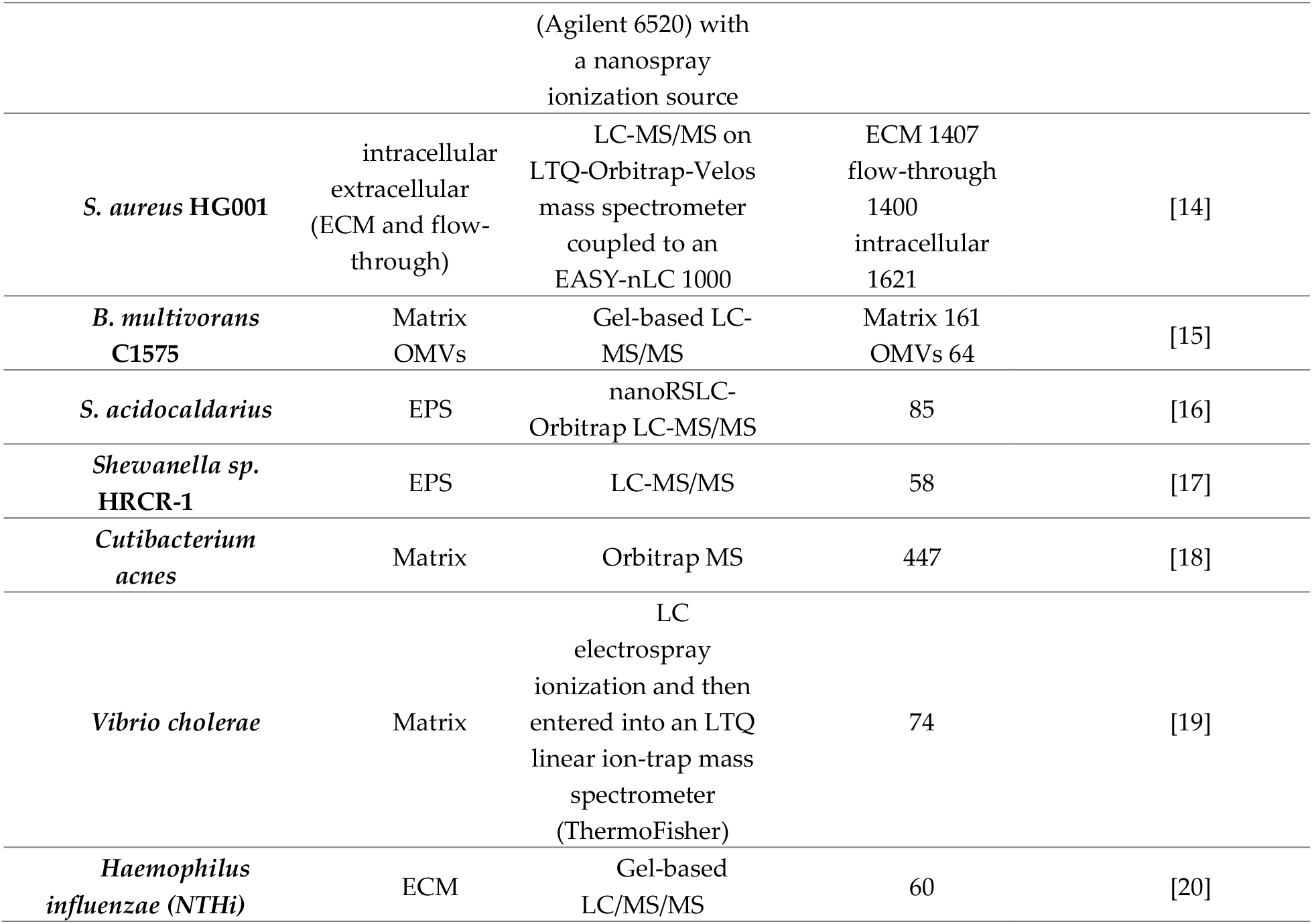
Studies devoted to biofilm matrix proteome. Biofilm’s compartments and MS methods were named by original publications.

Comparative study of liquid culture of PAO1 strain and clinical isolates showed that cystic fibrosis isolates expressed a narrower range of transporters and a broader set of enzymes of metabolic pathways for the biosynthesis of amino acids, carbohydrates, nucleotides, and polyamines, but this study did not cover biofilm mode of life as well as extracellular matrix composition [21]. Only one study described the proteome of the matrix in comparison with embedded cells [8] and one else study compared the matrix with the total biofilm proteome [7], both studies were devoted to reference strain PAO1. The gap in understanding the difference in protein composition between matrix and embedded cells frustrates the development of antibiofilm therapeutics and the overall understanding of biofilm biology. Here we performed a proteomic study of matrix composition in comparison with embedded cells for the clinical strain of *P. aeruginosa* to identify bioprocesses taking place in the matrix as probable targets or as factors or as barriers during pharmacological development of antibiofilm therapeutics.

## 2. Results

### 2.1. General overview of proteomes

*P. aeruginosa* KB6 (exoT+; exoY+; exoU-; exoS+; full name - GIMC5015:PAKB6) is a clinical isolate from bronchoalveolar lavage of patient from ICU [22]. This strain has strong biofilm-forming phenotype (Supplementary figure 2).

Two independent and separated in time biofilms were grown in a liquid medium. For the proteomic study, we have grown static biofilm for 18 h in LB medium. This time point corresponds to biofilm in the early stationary phase when the number of embedded bacterial cells reaches the plateau, and the matrix is already formed. Further increase in biofilm biomass occurs mainly due to extracellular components rather than an increasing number of bacterial cells. During the further biofilm growth, mass of extracellular substance significantly impacted with lysed cells. To decrease the number of proteins that may represent passive cell leakage or death during the biofilm stationary phase and to avoid interference of these «archeological» proteins with secreted extracellular proteins, we choose this early time point (18 h). To investigate protein composition of extracellular biofilm matrix, we used a previously published method of separation of biofilm matrix from embedded cells with high ionic solution of NaCl [23,24]. Extracellular matrix and embedded cells were separated and processed for protein isolation. Also, one more biological replicate (the third) representing embedded cells only was added, while matrix from this biofilm was used to check if protein quantity is enough for proteome analysis (total protein quantity from matrix preparation was more than 100 mkg, while minimum requirements is 50 mkg). We performed proteomic analysis for each sample in three technical replicates. For protein identification, *P. aeruginosa* KB6 strain-specific protein dataset was created based on genome sequence (available in GenBank under accession number NZ_CP034429). For protein identification, we used MSFragger software. The full list of all identified proteins is available in Table S1.

In total, we identified more than 1600 proteins in all samples. After initial manual inspection of LFQ intensities, we observed matrix-specific proteins, cell-specific proteins, and proteins with presence in both compartments. Spearman correlation coefficient for proteins LFQ intensities between cells and matrix was 0.4959 (95% CI 0,4555 to 0,5343) (Figure 1). While the correlation was expectedly positive, there were several examples of outfitters like lytic murein transglycosylase with well-known localization on the outer surface of the bacterial cell wall [25].

**Figure 1.**
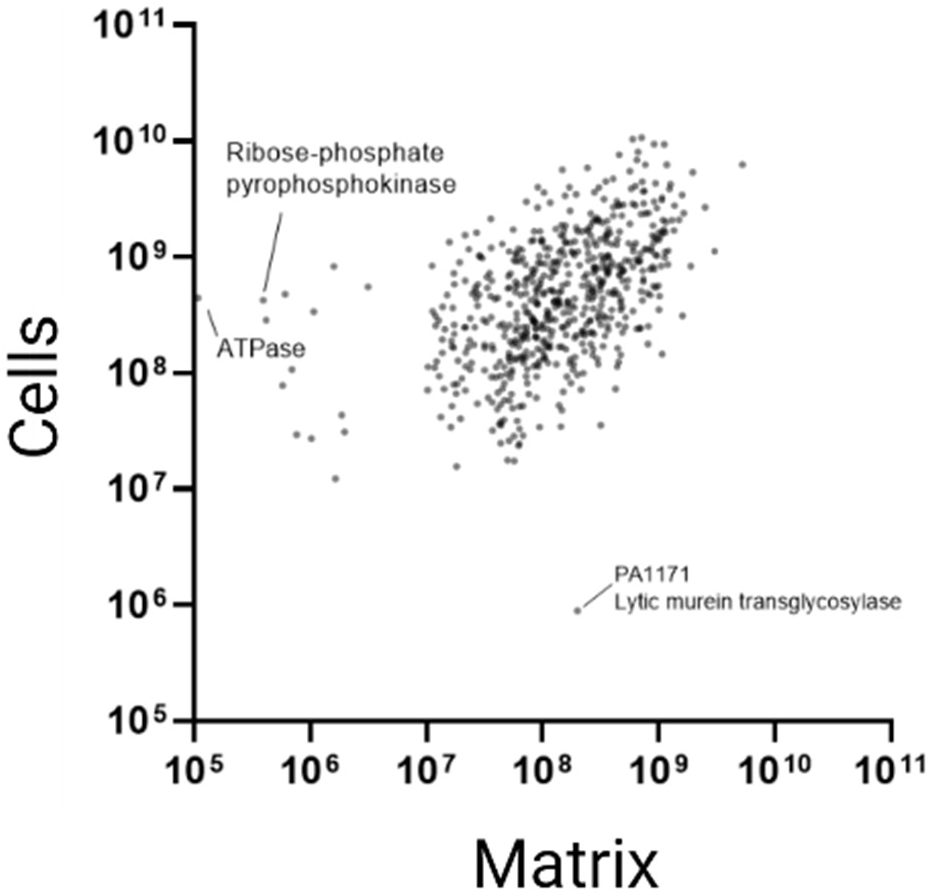
Correlation matrix of individual protein intensities (LFQ) between embedded cells and matrix. Proteins with zero LFQ intensities are out of axis range and not plotted.

While an inspection of intensity signals is not applicable for quantitative comparison due to different nature of cells and matrix samples, it is still giving us an additional consciousness about the proper separation matrix from cells. The absence of several intracellular proteins in matrix samples confirms that our approach to separate matrix from cells does not cause significant cell leakage during sample preparation. For example, proteins involved in ribosome assembly and function (L29, L33, S15, Era), septum formation and division (FtsX and MinC), transcription regulation (Cro/Cl family transcriptional regulator) and some others were absent in the matrix (LFQ intensities and spectral counts were zero in all matrix samples), while for some examples LFQ intensities in cells samples were more than 1*10^8^. So, the absence of these proteins in matrix samples confirms matrix separation without cell lysis.

Cells and matrix differ in overall biomolecule content, physicochemical and other properties. That was our premise for rigorous statistical analysis of comparative protein representation. For getting a quantitative comparison of the representation of proteins, we proceeded to logarithm transformation and quantile normalization of LFQ intensities. We made statistical analysis with the Limma R package [26]. All identified proteins are listed in table S1, and proteins differing in their representation in cells and matrix are listed in Table S2. For representation of the fold difference, volcano plot was created (Figure 2).

**Figure 2.**
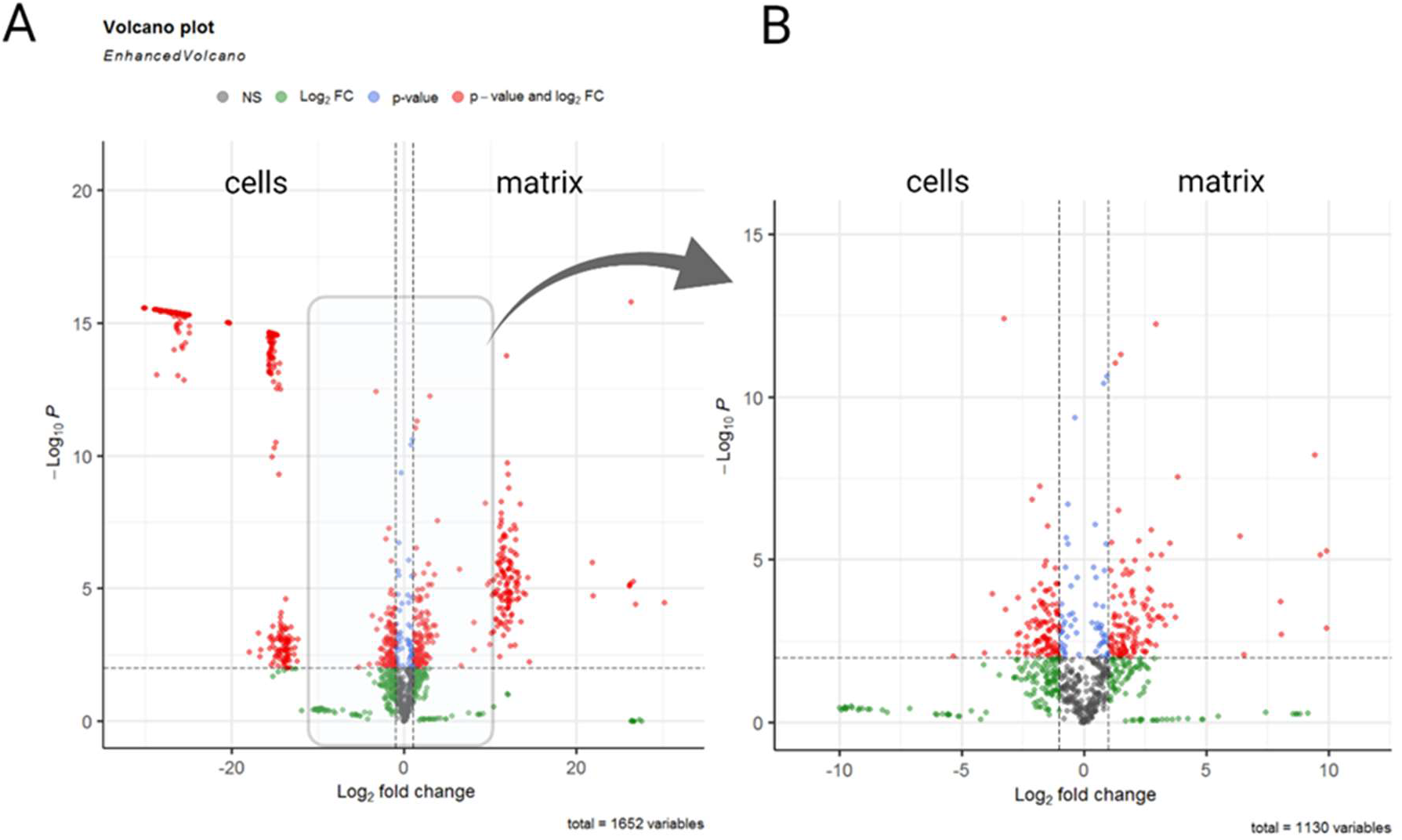
Volcano plot of representation of proteins between cells and matrix. A – overall protein representation; B – an enlarged area with proteins with fold change between Log2 from 1 to 10. Red color indicates proteins with fold change more than 2 and significance value p<0.01; green color indicates proteins with fold change more than 2 and significance value p>0.01; blue color indicates proteins with fold change less than 2 but significance value p<0.01; grey color indicates proteins with fold change less than 2 and significance value p>0.01.

In total, there were more than 1600 proteins, and we found 957 of them in the biofilm matrix (766 matrix proteins were present in both biological replicates).

### 2.2. Unique proteins in the extracellular matrix

Unique proteins were defined if we observed them (MaxLFQ total and unique intensities are non-zero) either in all matrix samples or in all cell’s samples. Totally ten proteins were present in the extracellular matrix only. We have proceeded with the literature search and manual biofilm-related functional annotation for these matrix unique proteins (Table 2). Nonetheless, for most extracellular matrix-only proteins, we did not find straightforward evidence of their importance for biofilm structure or any extracellular biofilm-related function.

**Table 2.**
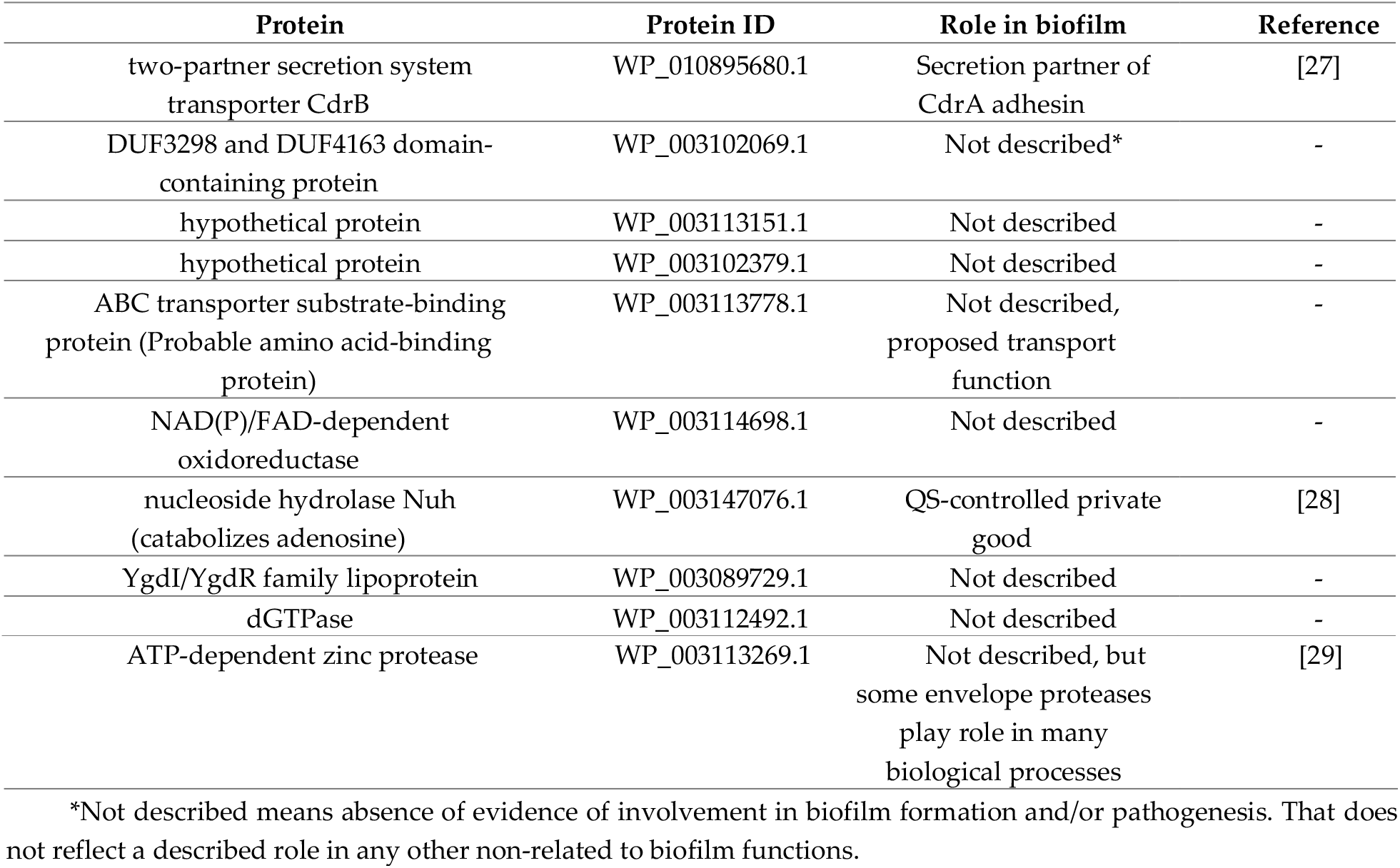
Matrix unique proteins. Matrix unique proteins had nonzero intensities in all matrix samples and zero intensities in all cell samples.

| Protein | Protein ID | Role in biofilm | Reference |
| --- | --- | --- | --- |
| two-partner secretion system transporter CdrB | WP_010895680.1 | Secretion partner of CdrA adhesin | [27] |
| DUF3298 and DUF4163 domain-containing protein | WP_003102069.1 | Not described* | - |
| hypothetical protein | WP_003113151.1 | Not described | - |
| hypothetical protein | WP_003102379.1 | Not described | - |
| ABC transporter substrate-binding protein (Probable amino acid-binding protein) | WP_003113778.1 | Not described, proposed transport function | - |
| NAD(P)/FAD-dependent oxidoreductase | WP_003114698.1 | Not described | - |
| nucleoside hydrolase Nuh (catabolizes adenosine) | WP_003147076.1 | QS-controlled private good | [28] |
| YgdI/YgdR family lipoprotein | WP_003089729.1 | Not described | - |
| dGTPase | WP_003112492.1 | Not described | - |
| ATP-dependent zinc protease | WP_003113269.1 | Not described, but some envelope proteases play role in many biological processes | [29] |

\*Not described means absence of evidence of involvement in biofilm formation and/or pathogenesis. That does not reflect a described role in any other non-related to biofilm functions.

Additionally, to unique matrix proteins, there were proteins with significant and extremely high log2 fold change (Log2FC) – more than 20. We considered these proteins as semi-unique matrix proteins (Table 3).

**Table 3.** Semi-unique matrix proteins. Semi-unique proteins had nonzero intensitiesLFQ in both cells and matrix samples and log2FC more than 20.

| Protein | Protein ID | Role in biofilm | Reference |
| --- | --- | --- | --- |
| malonate decarboxylase subunit alpha | WP_003112666.1 | Malonate improve biofilm formation | [30] |
| OprD family porin | WP_003114177.1 | Not described*, carbapenem binding | [31,32] |
| ABC transporter substrate-binding protein | WP_003098136.1 | Not described, proposed transport function | - |
| carbon-nitrogen hydrolase family protein | WP_003112739.1 | Not described | - |
| bifunctional riboflavin kinase/FAD synthetase | WP_003102619.1 | Not described; flavin nucleotide biosynthesis | - |
| M48 family metallopeptidase | WP_003112546.1 | Not described | - |
| transporter substrate-binding domain-containing protein | WP_003115039.1 | Not described, proposed transport function | - |
| division/cell wall cluster transcriptional repressor MraZ | WP_003103101.1 | Not described | - |

\*Not described means absence of evidence of involvement in biofilm formation and/or pathogenesis. That does not reflect a described role in any other non-related to biofilm functions.

### 2.3. Functional classification of proteins in the extracellular matrix

All overrepresented in matrix proteins are displayed in Figure 2A. We observed distinguishable clusters of overrepresented proteins depending on their fold change (Log2FC). While a separate cluster of highly overrepresented proteins with log2FC more than 10 is visible on the main volcano plot (Figure 2A), for better resolution of the area within log2FC 1-10 frame we also provide an enlarged area of the volcano plot (Figure 2B). Analysis of bioprocess classification of all found in matrix proteins and overrepresented in matrix proteins is displayed in Figure 3. Many bioprocess groups include at least one overrepresented in biofilm matrix protein.

**Figure 3.**
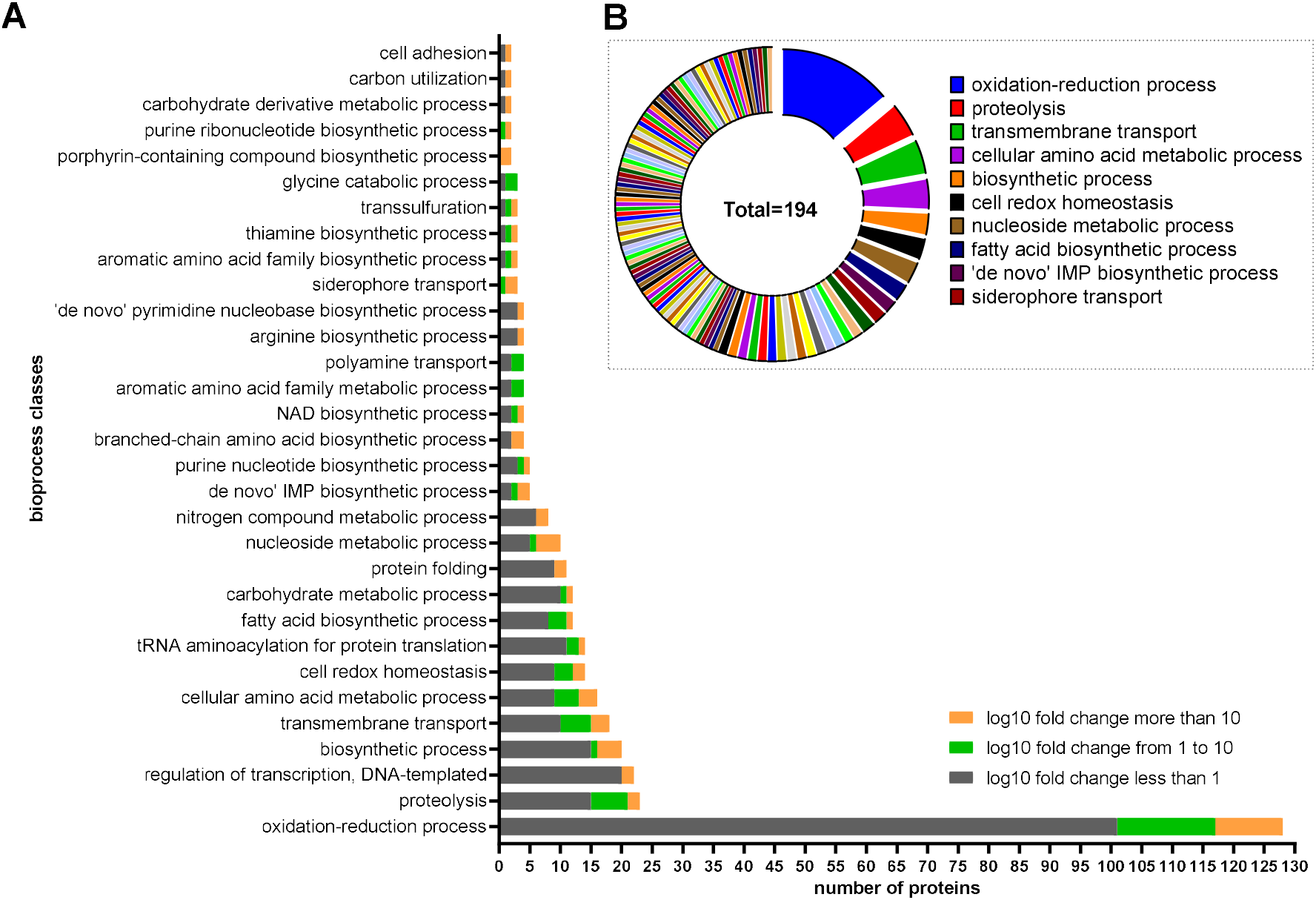
Bioprocess group classification of proteins found in the biofilm matrix. A – number of represented proteins in different bioprocess groups concerning log2 fold change in comparison with embedded cells: grey color – proteins found in the matrix but not overrepresented in comparison with embedded cells; green color – overrepresented proteins with log2 fold change from 1 to 10; orange color – overrepresented proteins with log2 fold change more than 10, only groups with at least 2 found in matrix proteins and at least one overrepresented protein are displayed; B – distribution of overrepresented proteins in biofilm matrix, each color represents individual bioprocess group, the legend indicates top ten of bioprocess groups.

We focused on groups with the largest number of found proteins and the proportion of overrepresented proteins from all found in matrix proteins. For this reason, we applied the following criteria: (1) at least 10 proteins per group found in the matrix; (2) more than 3 found in matrix proteins are overrepresented (4 to 28 overrepresented proteins from 8 bioprocess groups). Such bioprocesses belong to oxidation-reduction and cell redox homeostasis, proteolysis, transmembrane transport, amino acid, and nucleoside metabolic processes, fatty acids biosynthesis. An important consideration is that GO annotated bioprocesses consist of groups with highly different numbers of proteins. Number of proteins involved in bioprocess reflects complex nature and flexibility of protein interactions. As an absolute number of identified proteins may not reflect bioprocess representation, we count for each selected bioprocess group proportion of found proteins from all GO annotated group members as a measure of bioprocess representation (Table 4).

**Table 4.** Bioprocess representation in biofilm matrix based on the proportion of found proteins from GO-annotated for the bioprocess proteins.

|  | number<br>of found in<br>matrix<br>proteins | number of<br>overrepresented<br>in matrix proteins | GO-annotated<br>number of<br>proteins in<br>the database | %% of<br>found in matrix<br>proteins from<br>GO-annotated in<br>database |
| --- | --- | --- | --- | --- |
| oxidation-reduction process | 128 | 27 | 448 | 29 |
| proteolysis | 23 | 8 | 87 | 26 |
| biosynthetic process | 20 | 5 | 985 | 2 |
| transmembrane transport | 18 | 8 | 905 | 2 |
| cellular amino acid metabolic process | 16 | 7 | 141 | 11 |
| cell redox homeostasis | 14 | 5 | 26 | 54 |
| fatty acid biosynthetic process | 12 | 4 | 26 | 46 |
| nucleoside metabolic process | 10 | 5 | 11 | 91 |

In such analysis, the top three represented bioprocesses were cell redox homeostasis, nucleoside metabolism, and fatty acid synthesis. Considering the annotated number of proteins for each group, the nucleoside metabolic process was represented with ten proteins in the matrix out of eleven GO annotated proteins. In contrast to the nucleoside metabolic process, DNA-templated regulation of transcription (this group includes only 2 overrepresented in matrix proteins) and biosynthetic process groups were represented in matrix with a greater number of proteins (22 and 20, respectively), but it was less than 5% from GO annotated in database proteins for these groups (22 proteins from 472, and 20 from 985, respectively).

### 2.4. Matrix decreases cytotoxicity of bacteria against A549 lung epitelial cells

We hypothesized that the biofilm matrix may affect the way how bacteria interact with eucaryotic cells and form biofilm in coculture model. As some compounds may freely diffuse between matrix and surrounding medium, we served biofilm-conditioned medium for comparison. In experimental conditions suitable for eucaryotic cells (DMEM medium supplemented with fetal bovine serum (FBS), atmosphere of 5% CO2) addition of matrix to the isolated from biofilm bacteria improved biofilm formation while biofilm-conditioned LB medium slightly reduce biofilm biomass (Figure 4).

**Figure 4.**
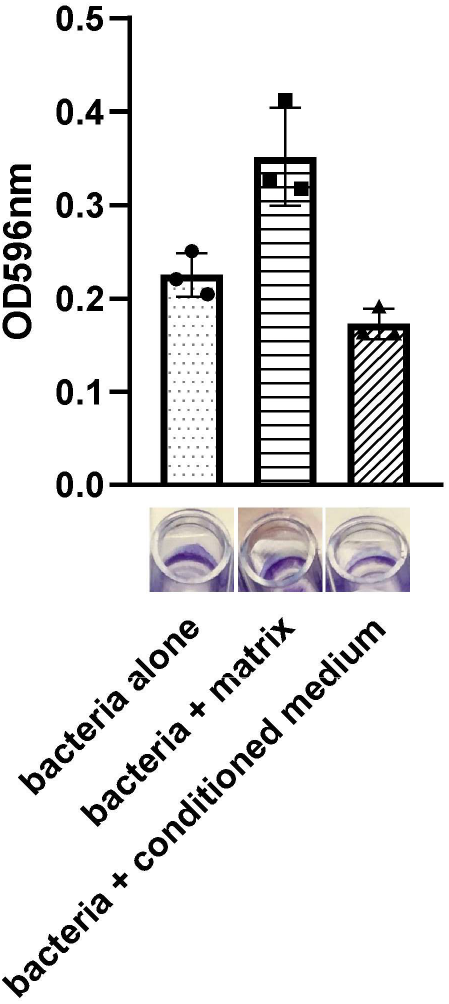
Matrix effect on biofilm formation at conditions suitable for eucaryotic cell culturing. Bacteria isolated from biofilm with addition of 10% of matrix or biofilm-conditioned medium were grown in DMEM medium supplemented with 10% FBS at 5% CO2 for 18 h, then stained with crystal violet. Diagram represents the quantification of CV staining and well photographs represent biofilm formation at the air-liquid interface.

For coculture experiments, we choose A549 adenocarcinomic human alveolar basal epithelial line as a pulmonary epithelial cell model. Addition of bacterial cells isolated from biofilm at MOI=3 resulted in A549 cell layer disruption and eucaryotic cell death after 18 h of incubation in 5% CO2 atmosphere. Addition of matrix to coculture model caused a significant change of A549 appearance with maintained eucaryotic cell attachment and monolayer integrity, while addition of biofilm-conditioned medium had an opposite action (Figure 5A).

**Figure 5.**
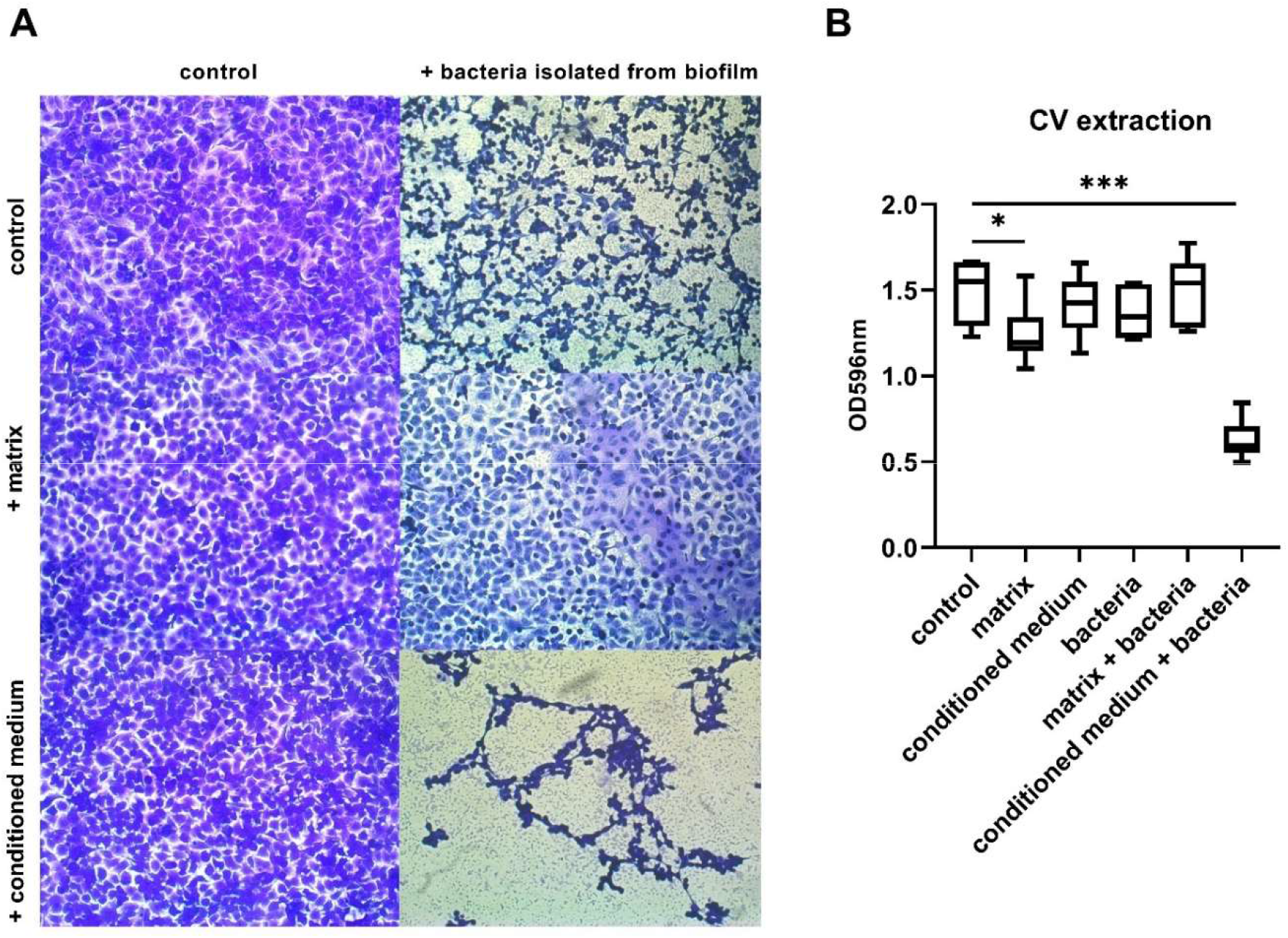
Matrix effect on biofilm formation and cytotoxicity in coculture isolated from biofilm bacteria with A549 eucaryotic cells. A – light microscopy at 10x magnification of coculture stained with crystal violet. B – cumulative biomass quantification with crystal violet. Asterisks indicate ANOVA p value: * - p<0.05; *** - p<0.001.

Importantly at this condition *P. aeruginosa* forms biofilm on both eukaryotic cells and plastic surface as well as pellicle biofilm on the air-liquid interface (Figures 4, 5 and Supplementary figure 2B), so quantification of staining showed cumulative biomass of eukaryotic cells and biofilm (Figure 5B). Bacteria alone disrupted monolayer of A549 cells, but some cells remained attached and biofilm formation on the walls of wells compensate biomass quantification. Matrix alone was slightly toxic to A549 cells but restricted bacterial cytotoxicity and the most A549 cells remained attached, so the overall biomass was as in the control. At the same time, well-tolerated biofilm-conditioned medium promotes bacterial cytotoxic effect, which is clearly visible under microscopy and after dye extraction. So the biofilm matrix was able to restrict bacterial cytotoxicity and prolong the maintenance of cell layer integrity.

## 3. Discussion

Biofilm Biofilm matrix in most cases contains many proteins. Extracellular proteins in the biofilm matrix provide their functions for the whole bacterial community, so they may be considered as public goods. Extracellular functions of matrix proteins include (but are not limited to) the external digestive system, signaling, protection, and maintaining the stability of the matrix [1,33–35]. Moreover, many bacterial proteins may have moonlight functions [36]. So studies devoted to matrix composition including proteomics are essential for depicting biofilm lifestyle.

We showed that the matrix may suppress bacterial cytotoxicity in coculture with eukaryotic cells. Biofilm-associated infections frequently cause local tissue damage and inflammation without system dissemination. While this complex process includes counteraction of host immune system and virulence of bacterial pathogen, biofilm matrix might play a direct role in limitation of barrier tissue injury and localization of infection process. There are many ways how matrix could restrict bacterial «aggression». Just a few of them include providing favorable nutrients rather than eukaryotic cells, restriction of bacterial sensing of eukaryotic cells, suppression of virulence activation and others. Wang et al. have reported that cytotoxicity is attenuated at high MOI in coculture of *P. aeruginosa* with A549 cells [37]. Authors identified phenylacetic acid (PAA) as compound responsible for T3SS suppression. PAA was identified in the conditioned culture medium from planktonic bacterial culture. In contrast, we did not observe any protective role of biofilm-conditioned medium, but the idea and overall principal of virulence suppression by bacterial population in the stationary phase is similar. Kaya et al. reported 90% survival of PBMC co-cultured with *P. aeruginosa* biofilm [38]. Moreover, PBMC responded to *P. aeruginosa* biofilm and vice versa - biofilm increased bacterial cells number in the presence of PBMC or PBMC-secreted factors. This observation indicates the existence of cross-talk during establishing of chronic infection and our data suggest matrix importance in this communication.

Understanding the protein composition of the matrix is critical for resolving both fundamental questions in bacterial lifestyle and the development of tools for manipulating biofilms. Most proteomic studies focus on the whole biofilm and usually compare the whole biofilm proteome with the planktonic cells proteome. At the same time, the identification of matrix protein composition may open new opportunities for the development of antibiofilm drugs. Well known that biofilms of pathogenic microorganisms are tolerant to antibiotics, many other therapeutics, and host immune factors due to the matrix [39]. Matrix-disrupting or interrupting agents may reverse tolerant phenotype and increase the efficacy of antibiotic therapy [40]. For example, antibody-mediated destabilization of the matrix through the disruption of IHF-DNA complexes was effective in vitro and in vivo as a single therapy and in combination with antibiotics [41–43]. Moreover, extracellular targets less probable cause selection of resistant mutants, so targeting of matrix proteins is a perspective way to combat chronic infections, especially infections caused by ESKAPE pathogens. Despite all of these, it is still little known about matrix proteomes. Only numerous studies have described the matrix proteome of reference strain PAO1 of *P. aeruginosa* and some other bacteria (Table 1). Here we for the first time describe the matrix proteome of clinical isolate of *P. aeruginosa* in comparison with embedded cells. In comparison with other studies, we identified the largest number of proteins in the matrix. While some proteins might be invisible due to their low concentrations and individual limits of detection, we believe that further improvement of MS equipment and techniques will get a more comprehensive picture of bacterial proteomics.

In our study, only a small number of proteins were unique for matrix. One of them – CdrB protein is involved in the transport of CdrA adhesin. This adhesin is important for biofilm formation and its binding to Psl results in increased biofilm structural stability. Antibody-mediated blocking of CdrA inhibit biofilm formation[44]. Also, *cdrAB* is regulated together with Psl. CdrA binding to Psl protects it from endogenous and exogenous protease digestion [45]. Both CdrA and Psl coding operons are present in the genome of KB6. Surprisingly, MaxLFQ unique intensities for CdrA protein were zero in both cells and matrix samples. In a less sensitive gel-based proteomic studies, authors have observed CdrA protein in the matrix [7, 20], but in our data CdrA seems to be under the limit of quantitation. One of the possible explanations is that endogenous proteases had degraded CdrA by the time when we collect biofilm and other mechanisms maintained biofilm structure. In this case found in matrix CdrB might be «archeological» protein. Further dynamic studies in Psl and CdrA presence in the matrix may shed a light on this question.

Another matrix unique protein is nucleoside hydrolase (Nuh) – an enzyme that hydrolyzes adenosine and inosine, allowing the cell to grow on these nucleosides as the sole carbon or nitrogen source. Nuh was considered as an intracellular (periplasmic) private good [28]. In our study, we found Nuh in the matrix, but not inside bacterial cells. That means Nuh might be an extracellular public good, at least for some strains like ours. Also, transporters responsible for adenosine transport to periplasmic space in *P. aeruginosa* remain still undiscovered, so if Nuh works outside the cell, the need of transporters is questionable. In an environment with adenosine as the sole carbon source, Nuh mutant has impaired growth [46], but therapeutic potential of direct or indirect inhibition of Nuh activity remains elusive due to the nutrient-rich nature of infected tissues, and further research is needed.

For the rest 8 matrix-unique proteins, there is no clear evidence of their possible role in biofilm. Meanwhile, as matrix is considered a nutrient-rich environment, presence of substrate-binding protein from ABC transporter and its role in nutrient (probably amino acids) acquisition is obvious. Antibody-mediated blocking of some ABC transporters was shown to be effective in vitro and in vivo against *M. hominis* and *S. aureus* [47,48]. Monoclonal antibody Aurograb® entered phase III clinical trial as an addition to vancomycin therapy for deep-seated staphylococcal infections (NCT00217841), but the trial was stopped due to lack of reaching the primary endpoint. Anyway, somewhere positive results in targeting eukaryotic ABC transporters for cancer treatment support the idea of a broader evaluation of the similar capability for prokaryotes. Also, transmembrane transport is one of the prevalent bioprocess groups in terms of all found in matrix proteins (n=18) as well as in terms of several overrepresented in matrix protein (n=8).

ATP-dependent Zn proteases are common enzymes in the cell envelope of *P. aeruginosa*, they participate in several processes, including metabolism, protein transport and removal of misfolded proteins, and adaptation to environmental conditions [29]. Also, some proteases may act as a virulence factor - Zn2+-dependent protease *Bacillus anthracis* called Lethal Factor is required for infection [49]. The proteolysis bioprocess group was the second represented in the matrix with 8 overrepresented proteins. Proteolysis is a part of the external digestive system and provides peptides and amino acids for bacterial nutrition, so the presence of some proteins involved in amino acids metabolism was expected (this bioprocess group includes 16 found in matrix proteins with 7 overrepresented proteins). Proteolytic activity could be crucial for both bacterial survival and infection process and targeting bacterial proteases could be a perspective way to combat bacterial infections [50]. Moreover, for Zn proteases, host nutritional immunity (including Zn-dependent processes) was effective against infections caused by *P. aeruginosa* [51]. Also, deprivation of Zn ions was proposed to combat infections caused by another common pathogen - S. aureus [52].

Despite a small number of matrix unique proteins, we found a lot of overrepresented proteins in the matrix. Eight proteins with extremely high log2 fold change (more than 20) were considered as semi-unique for matrix, but we did not find in the literature any role in biofilm lifestyle. So as for unique matrix proteins, there is an unexplored area in biofilm biology.

Bioprocesses classification of represented in matrix proteins reveals several groups. Considering log2 fold change, we found that groups of oxidation-reduction processes, biosynthetic processes, and nucleoside metabolism had the largest number of highly overrepresented proteins. Obviously, each bioprocess might vary in the number of involved proteins, so we also introduced bioprocess representation as a part of all GO-bioprocess annotated proteins.

In the matrix, the most reach group of proteins (128 proteins found in the matrix, 27 overrepresented) belongs to oxidation-reduction processes. Also, a group of proteins involved in cell redox homeostasis was one of the most represented GO-annotated bioprocesses in the matrix (14 proteins found, 8 overrepresented from 26 annotated in GO bioprocess database). Biofilms of *P. aeruginosa* contain molecules involved in virulence and competition with other microorganisms, including redox-active molecules. Self-produced factors involved in the generation of reactive oxygen species might be harmful to the internal bacterial community. So *P. aeruginosa* is balancing to maintain oxidation-reduction processes at the appropriate level, so the balance of oxidation-reduction reactions and redox homeostasis likely play a significant role in the biofilm matrix as an environment with limited diffusion. Several effective therapeutic approaches utilize oxidative stress to combat bacterial biofilms, including photodynamic therapy (PDT) and sanitizers like hydrogen peroxide [53,54]. Also, extracellular electron transfer (EET) exists inside biofilm matrix, but the role of the protein component of EET remains undiscovered [55].

The second represented bioprocess included proteins involved in the fatty acid biosynthetic process (4 overrepresented proteins from 12 found in the matrix). Fatty acids are one of the major components of the cell envelope. Also, it is a well-known signal function of cis-2-docenoic acid messenger (DSM) as well as the importance of the fatty acid component of AHL [56]. In a recently published study, Altay et al. made a comprehensive analysis of essential reactions and affected pathways in *B. cenocepacia* (both planktonic and biofilm) using a systems biology approach. From all identified essential reactions, lipid metabolism was responsible for more than half of the single lethal reactions; among this fatty acid biosynthesis was most frequently found [57]. That data supports the further development of fatty acids metabolism inhibitors as promising therapeutics against bacterial infections, including bacterial biofilms.

The most represented bioprocess was nucleoside metabolism – 10 out of 11 GO-annotated proteins were found in the matrix. Biofilms are often enriched with extracellular nucleic acids, which act not only as a structural component or component of EET, but also as a nutrient source [58]. Nucleosides act as substrates and cofactors in many biosynthetic processes, as signal molecules, and are involved in regulating bacterial community inside biofilm [59], so the presence of nucleoside metabolic proteins is required. Therapeutic targeting of proteins involved in these processes is theoretically possible, but at present is not clear. At the same time, nucleoside analogs are common drugs in other nonbacterial diseases, and evaluation of their possible role as antibacterial drugs may open new opportunities [60]. So antimycotic drug 5-fluorocytosine was able to suppress virulence of *P. aeruginosa* in a murine model of lung infection [61].

Obviously, sources of extracellular matrix proteins belong to two main categories: (1) active secretion of biomolecules and (2) passive way to increase extracellular content as a consequence of cell leakage or lysis. While some extracellular proteins are «passive» products of bacterial cell lysis, they still might be active outside the cell and provide their function to the bacterial community extracellularly. As protein degradation rates lay in broad ranges, some proteins may present in the matrix for a long time after leakage or secretion from the cell, so their occurrence in the matrix does not (1) match the actual situation inside cells or (if. e. active transcription and translation), or (2) reflect any real extracellular needs for biofilm (i.e. be structurally or/and physiologically involved in extracellular processes). The one limitation of our study is an inability to conclude if the protein is «archeological», bystander, or functionally active in the matrix and how these proteins are distributed in the matrix (are they cell-attached, part of OMVs, or associated with other matrix components). Moreover, in a prism of drug development, some proteins may act as distracting extracellular targets in a way of absorption of active drug and distract from intracellular targets. Also there is a risk of potentiating severe infection in a case of matrix disruption and massive release of bacterial cells. To choose a right strategy for antibiofilm development, there is a need for detailed knowledge about function and dynamic of every single protein. Another important limitation is the fact that biofilm cultured *in vitro* on the plastic surface in bacteriological mediums does not reflect real physiological conditions and extensive study must be done to evaluate the relevance of any *in vitro* results for the understanding of pathogenesis of chronic infections and finding targets for antibiofilm drug development.

## 4. Materials and Methods

### 4.1. Biofilm growth and separation matrix from cells

*P. aeruginosa* KB6 (clinical isolate) was a gift from Zigangirova N. A. (Gamaleya NRCEM, Moscow, Russia) [21]. For biofilm preparation single colony from TSA plate was picked in liquid LB medium and grown 24 h at 37 C, 210 rpm. The liquid culture was diluted 50 times with LB medium in a volume of 20 ml and placed in Petri dishes for 18 h at 37 C under static conditions. Then the medium was removed, and biofilm was exposed to 10 ml of 1.8 M NaCl. After 5 min of incubation bacterial suspension and dissolved matrix were separated with centrifugation at 5000 g. Liquid phase (dissolved matrix) was filtered through a 0.22 mkm syringe filter. Protein was precipitated with cold acetone (up to 80 %) 18 h at −20 C. Cells pellet was resuspended in lysis buffer (2% SDS; 50 mM Tris-HCl; 180 mM NaCl; 0,1 mM EDTA; 1 mM MgCl2) and boiled for 15 min in a water bath. Cell debris was removed by centrifugation at 10000 g, 15 min, and proteins from the liquid phase were precipitated with 80% cold acetone as for matrix samples. Precipitated proteins were pelleted with centrifugation at 10000 g, 20 min, 4 C. Pellet was washed two times with 80 % cold acetone and proceeded for proteomic sample preparation. Total protein quantity was measured with QuDye Protein kit (Lumiprobe) on Qubit fluorometer (ThermoScientific).

### 4.2. Proteomic sample preparation and peptide identification

Proteomic sample preparation and peptide identification were made in Advanced Mass Spectrometry Core Facility (Skolkovo Innovation Center, Moscow, Russia). The protein pellet was subjected to tryptic in-solution digestion. LC-MS/MS was carried out on a Q Exactive HF (Thermo Scientific) with a nanoESI interface in conjunction with an Ultimate 3000 RSLC nano HPLC (Dionex Ultimate 3000). Peptides were loaded onto a trap column and separated on an analytical column (C18) using an H2O/acetonitrile gradient with 0.1% formic acid for 150 min. The Q Exactive HF spectrometer was operated in the data-dependent mode with a nanoESI spray voltage of 1.8 kV, capillary temperature of 210 °C, and S-lens RF value of 55%. All spectra were acquired in positive mode with full scan MS spectra scanning from m/z 310–1500 in the FT mode at 120,000 resolution. A lock mass of 445.120025 was used. The top 25 most intense precursors were subjected to rapid collision induced dissociation (rCID). Dynamic exclusion with of 70 seconds was applied for repeated precursors.

### 4.3. Data analysis

To identify and quantify tryptic peptides and the proteins from which the peptides are derived, spectra from the MS/MS experiments were analyzed by GUI FragPipe v. 17.1 (https://github.com/Nesvilab/FragPipe). Peptide identification was performed by MSFragger search engine [62,63] using protein sequence database extracted from NCBI *(Pseudomonas aeruginosa* strain GIMC5015:PAKB6 chromosome, complete genome NZ_CP034429) with decoys and contaminants. Oxidation of methionine and acetylation of protein N-termini were set as variable modifications, carbamidomethylation of cysteine was set as a fixed modification. The maximum allowed variable modifications per peptide was set to 3, mass tolerance was set as 20 ppm for precursor and 0.02 Da for fragment ions. Philosopher kit tools [64,65] were used to estimate identification FDR. The PSMs were filtered at 1% PSM and 1% protein identification FDR. Quantification by label-free protein quantitation method and MBR was performed with IonQuant [66]. Obtained quantified data (intensities) were processed for differential expression analyses with limma package R [26], with followed visualization result by EnhancedVolcano package R (“EnhancedVolcano: Publication-ready volcano plots with enhanced coloring and labeling.” https://github.com/kevinblighe/EnhancedVolcano).

Bioprocess classification and functional annotation were made with the Pseudomonas Genome Database (https://pseudomonas.com/primarySequenceFeature/list?strain_ids=10430&term=Pseudomonas+aeruginosa+GIMC5015%3APAKB6&c1=name&v1=&e1=1&assembly=complete) [67].

For correlation analysis, ANOVA comparison and graphs, we used GraphPad Prism 9 desktop software (version 9.2.0). For the graphical abstract and figures arrangement we used bio-render online software (Biorender.com).

### 4.4. Coculture bacterial cells with eukaryotic cell line A549

Biofilm-conditioned medium, bacterial cells and dissolved matrix were prepared as described in section 4.1. Conditioned medium and matrix were filtered through a 0.22 mkm syringe filter. Bacterial cells pellet was resuspended in DMEM medium (Gibsco), and tenfold serial dilutions was plated on LB agar to count bacterial number. Samples were stored on ice and used within several hours after preparation. Adenocarcinomic human alveolar basal epithelial A549 cell line was maintained in DMEM medium (Gibsco) supplemented with 10% fetal bovine serum (Gibsco). A549 cells were seeded in 96-well tissue culture plate at 2*10^4^ cells per well and cultured to reach 85 % confluence. Then 10% of medium volume in well were replaced with matrix, biofilm conditioned medium and bacterial cells suspension in desired combinations. For the control well we used 1.8 M NaCl diluted as in matrix preparation and DMEM medium as for bacterial dilution. Plates were placed in atmosphere of 5% CO2, 37 ^o^C for 18 h. After incubation liquid was removed from the plate, wells were washed with 0.9% NaCl, fixed with ice cold methanol and stained with crystal violet (BD) for 30 minutes. After washing with tap water plates proceeded to light microscopy. For quantification crystal violet dye was extracted with 30% acetic acid and OD495nm was measured on plate spectrophotometer (ThermoScientific).

## 5. Conclusions

Biofilm matrix of clinical strain P. aeruginosa contains hundreds of proteins. There are several unique for matrix and many overrepresented in matrix proteins, which reflect several bioprocesses. Development of antibiofilm therapeutics may benefit in the case of targeting proteins and processes taking place in the biofilm matrix as the sole mechanism of action or in combination with antibiotics. Altogether, matrix protein composition is important for choosing a successful strategy in antibacterial drug development and reaching unmet needs of curing biofilm infections.

## Supplementary Materials

The following supporting information can be downloaded:

Table S1: All identified proteins.

Table S2: Differentially represented proteins.

Supplementary Figure S1: Number of recovered CFU after matrix separation

Supplementary Figure S2: Biofilms of *P. aeruginosa* KB6 stained with crystal violet

## Author Contributions

Conceptualization, D.E., A.S., N.P.; methodology, A.S., K.D.; software, A.S., N.P.; validation, A.S.; investigation, D.E., I.K., V.R.; resources, Y.R.; data curation, M.D.; writing–original draft preparation, D.E., M.D., I.T.; writing–review and editing, Y.R.; project administration, D.E.; funding acquisition, A.G. All authors have read and agreed to the published version of the manuscript.

## Data Availability Statement

The mass spectrometry proteomics data have been deposited to the ProteomeXchange Consortium via the PRIDE [62] partner repository with the dataset identifier PXD031480.

## Acknowledgments

We thank Nailia A. Zigangirova (Gamaleya NRCEM, Moscow, Russia) for the *P. aeruginosa* KB6 strain. We thank Viktor G. Zgoda from Advanced Mass Spectrometry Core Facility (Skolkovo Innovation Center, Moscow, Russia) for technical consulting and MS data obtaining.

---

**Figure S1.**
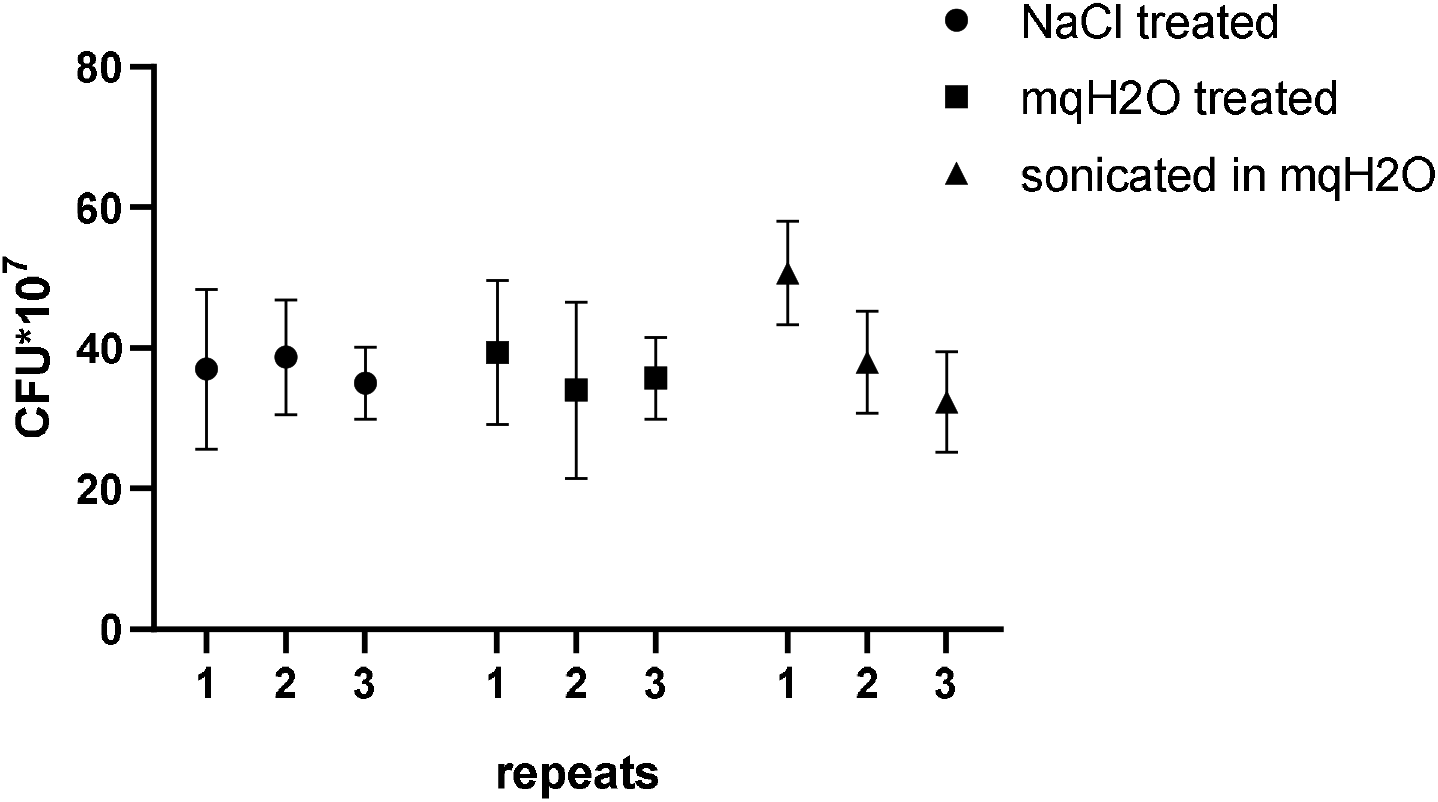
Number of recovered CFU after matrix separation. Biofilms in 96-well plate were separated with NaCl, intensive resuspension in mqH2O or sonication in US bath for 5 min. Graphs represent three replicated wells. Bacteria from each well were seeded in triplicates. Number of recovered CFU was calculated from number of CFU in last dilution with more than 20 CFU per plate.

**Figure S2.**
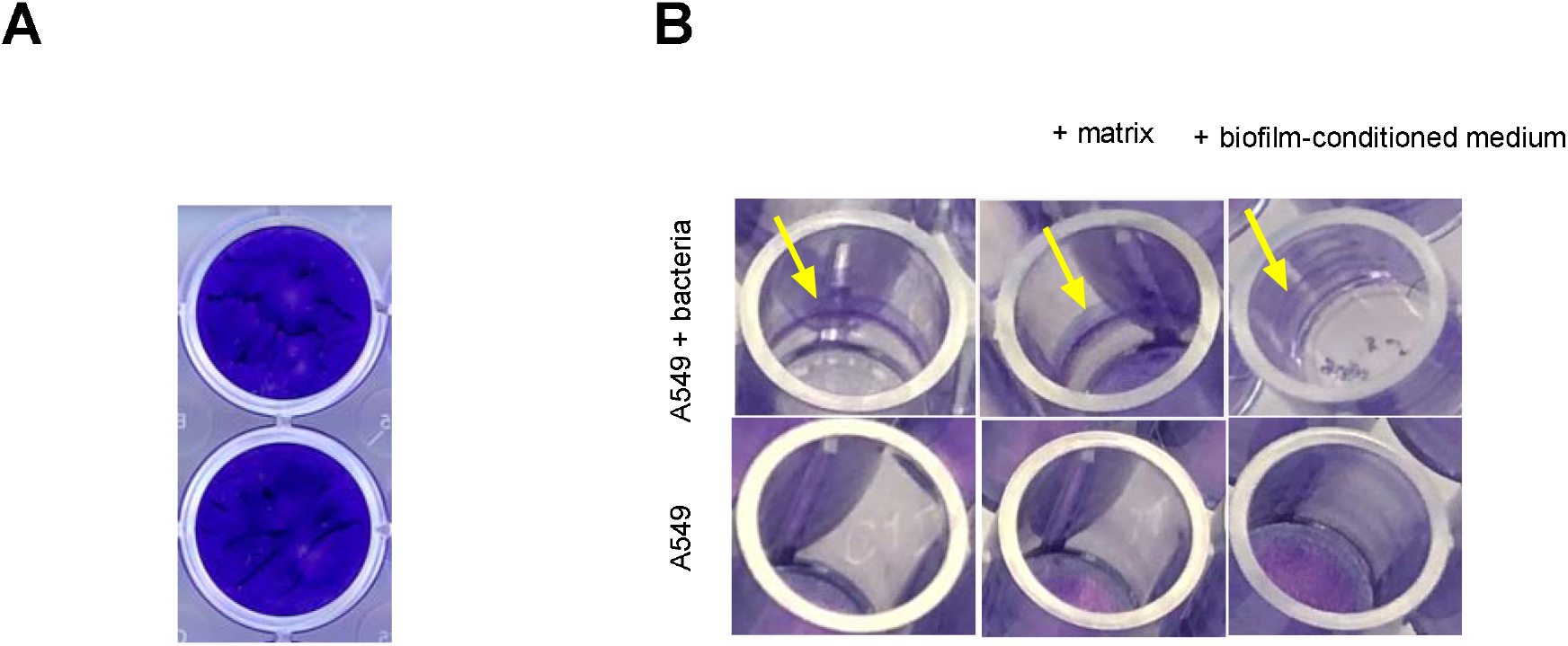
Biofilms of *P. aeruginosa* KB6 stained with crystal violet. **A** – KB6 demonstrates strong biofilm-forming phenotype. Crystal violet (CV) staining of biofilm formed in LB medium, 37 °C, 18 h. **B** – bacterial cells isolated from biofilm form biofilm in coculture model with A549 eukaryotic cells, MOI=3, 5% CO2, 37 °C, 18 h. Photographs demonstrate formation of biofilm at air-liquid interface (also biofilms are present at the bottom of wells in close proximity and in direct contact with A549 cells), so biofilm biomass has a significant impact in quantification of CV staining in coculture experiments.

